# Toddlers later diagnosed with autism exhibit multiple structural abnormalities in temporal corpus callosum fibers

**DOI:** 10.1101/099937

**Authors:** Noa Fingher, Ilan Dinstein, Michal Ben-Shachar, Shlomi Haar, Anders M. Dale, Lisa Eyler, Karen Pierce, Eric Courchesne

## Abstract

Interhemispheric functional connectivity abnormalities are often reported in autism and it is thus not surprising that structural defects of the corpus callosum (CC) are consistently found using both traditional MRI and DTI techniques. Past DTI studies however, have subdivided the CC into 2 or 3 segments without regard for where fibers may project to within the cortex, thus placing limitations on our ability to understand the nature, timing and neurobehavioral impact of early CC abnormalities in autism. Leveraging a unique cohort of 97 toddlers (68 autism; 29 typical) we utilized a novel technique that identified seven CC tracts according to their cortical projections. Results revealed that younger (<2.5 years old), but not older toddlers with autism exhibited abnormally low mean, radial, and axial diffusivity values in the CC tracts connecting the occipital lobes and the temporal lobes. Fractional anisotropy and the cross sectional area of the temporal CC tract were significantly larger in young toddlers with autism. These findings indicate that water diffusion is more restricted and unidirectional in the temporal CC tract of young toddlers who develop autism. Such results may be explained by a potential overabundance of small caliber axons generated by excessive prenatal neural proliferation as proposed by previous genetic, animal model, and postmortem studies of autism. Furthermore, early diffusion measures in the temporal CC tract of the young toddlers were correlated with outcome measures of autism severity at later ages. These findings regarding the potential nature, timing, and location of early CC abnormalities in autism add to accumulating evidence, which suggests that altered inter-hemispheric connectivity, particularly across the temporal lobes, is a hallmark of the disorder.

## Introduction

A leading hypothesis regarding autism neurophysiology suggests that the disorder is characterized by atypical anatomical and functional connectivity. This hypothesis has been supported by numerous Diffusion Tensor Imaging (DTI) studies that have assessed the microstructure of white matter fibers (Barnea-Goraly et al. 2004; Alexander et al. 2007; Ben Bashat et al. 2007; Thomas et al. 2011; Travers et al. 2012; Wolff et al. 2012; Ameis and Catani 2015; Solso et al. 2016) and by functional magnetic resonance imaging (fMRI) studies that have assessed functional synchronization across brain areas during different tasks (e.g., Just, Cherkassky, Keller, & Minshew, 2004; Müller et al., 2011) as well as during rest or sleep (e.g., Anderson et al., 2011; Di Martino et al., 2014; Dinstein et al., 2011).

Among the different forms of connectivity, inter-hemispheric anatomical and functional connectivity is of particular interest to autism research for several reasons. First, anatomical studies have reported that one or more subregions of the corpus callosum (CC) are smaller in older children, adolescents, and adults with autism (Egaas et al. 1995; Frazier and Hardan 2009; Travers et al. 2015, but also see Haar et al. 2014). Second, fMRI studies have reported that toddlers, children, adolescents, and adults with autism exhibit decreased inter-hemispheric functional connectivity during rest/sleep in comparison to controls (e.g., Anderson et al., 2011; Dinstein et al., 2011; Di Martino et al., 2014). Third, DTI studies have reported that adolescents and adults with autism exhibit smaller CC volumes, reduced fractional anisotropy (FA) and increased Mean Diffusivity (MD) values in comparison to controls (Alexander et al. 2007; Thomas et al. 2011; Catani et al. 2016). Fourth, up to 45% of children who are born without a CC (agenesis of the CC) exhibit behavioral symptoms that are consistent with a formal autism diagnosis (Lau et al., 2013; Paul et al., 2014). Similarly, BTBR mice, a strain where the corpus callosum is entirely absent (Wahlsten et al. 2003), exhibit several autism-like behaviors (Bolivar et al. 2007) and are a popular animal model for autism research (McFarlane et al. 2008; Silverman et al. 2010; Blanchard et al. 2012). Finally, studies of affect demonstrate that the integrity of the CC is important for higher-order social cognitive tasks (Symington et al. 2010; Mike et al. 2013), a key area of impairment in autism. Taken together, it is tempting to speculate that alterations in inter-hemispheric connectivity may represent an important neural phenotype of at least some individuals who develop autism.

How early and where exactly do interhemispheric connectivity abnormalities emerge? Since autism is a disorder of early neural development (Courchesne et al., 2011; Willsey et al., 2013; Stoner et al., 2014), it is particularly important to examine inter-hemispheric connectivity at very young ages when the behavioral symptoms of autism first emerge (Courchesne et al. 2007; Pierce et al. 2011). Only a few DTI studies have done so and all have reported that the CC at young ages in autism exhibits abnormally *increased* FA values (Ben Bashat et al. 2007; Weinstein et al. 2011; Xiao et al. 2014; Travers et al. 2015; Solso et al. 2016). This finding stands in sharp contrast to findings in mature children, adolescents and adults where nearly every study has reported *reduced* CC FA values in autism (Barnea-Goraly et al. 2004; Alexander et al. 2007; Jou et al. 2011; Travers et al. 2012; Vogan et al. 2016). Studies have suggested that the FA values transition from abnormally high to abnormally low values very early in development, apparently sometime between the ages of 2 and 4 years (Ben Bashat et al. 2007; Weinstein et al. 2011; Travers et al. 2015; Solso et al. 2016). These findings are in line with the transient early overgrowth hypothesis, which suggests that some infants and toddlers with autism have excess cortical neurons and display early accelerated brain growth followed by later arrested neuronal growth and axonal and synaptic development (Courchesne et al. 2001; Courchesne, Campbell, et al. 2011; Courchesne, Mouton, et al. 2011; Chow et al. 2012).

All of the DTI studies performed to date in toddlers with autism have subdivided the CC roughly into two or three segments without determining the cortical projections of the fibers in each segment. Thus, each segment likely contained a mixture of fibers that projected to multiple cortical areas. A more detailed examination of CC development, which takes into account the projection of CC fibers into specific occipital, parietal, temporal, and frontal areas is, therefore, important for determining which inter-hemispheric connections develop abnormally in autism and for revealing the nature and timing of abnormalities in each fiber group.

In the present study of prospectively identified toddlers with autism, we examined diffusion properties in seven different CC segments that were defined according to the cortical regions that were connected by their corresponding fibers. This allowed us to separate tracts that connected occipital, parietal, temporal, and frontal regions and sub-regions in each participant. We then examined the diffusion properties within a 10mm mid-sagittal segment of each tract, (i.e., the segment where tracts cross the mid-line) and compared the results across autism and control toddlers who were 1 to 4 years old at the time of MRI scanning. An important advantage of our intentional focus on midsagittal CC segments is that these segments do not contain crossing or kissing fibers (Jeurissen et al. 2013), which are known to alter diffusion measures (Mori and van Zijl 2002; Assaf and Pasternak 2008). This means that potential differences in diffusion measures across groups are more likely to represent true differences in underlying axonal microstructure rather than differences in the number of crossing fibers.

## Materials and methods

### Subjects and Recruitment

Toddlers were recruited through community referral and a population based screening approach called the 1-Year Well Baby Check-Up Approach (Pierce et al. 2011) at the University of California, San Diego. Ninety-seven toddlers participated in this study: sixty-eight with autism (mean age: 31 months old, range: 13 to 51 months) and twenty-nine typically developing controls (mean age: 29 months old, range: 13 to 48). There was no significant group difference in age (p > 0.05, two-tailed t-test for independent samples with unequal variance). All toddlers were scanned late at night, during natural sleep without the use of sedation (Pierce 2011). Parents of toddlers provided written informed consent and were paid for their participation. The UCSD human subject research protection program approved all experimental procedures. Diffusion MRI data from 14 of the 29 control toddlers and 25 of the 68 toddlers with autism was included in previous analyses reported by Solso et al. (2015).

### Diagnosis and behavioral evaluations

Diagnostic evaluations were performed by an experienced licensed clinical psychologist and were based on the Autism Diagnostic Observation Schedule (ADOS) (Lord et al. 2000) and clinical judgment. Additional evaluations included the Vineland Adaptive Behavior Scales, second edition (Sparrow 2011), which measure communication, daily living skills, socialization, and motor skill capabilities, as well as the Mullen Scales of Early Learning (Mullen 1995), which measure expressive language, receptive language, gross motor, fine motor, and visual reception capabilities (Table 1). Over 90% of the toddlers in both groups completed all three tests within 12 weeks of the MRI scan. In the current sample, 18 toddlers with an eventual diagnosis of autism who were 26 months old or younger at the time of initial diagnostic evaluation (and scan). All of these toddlers participated in a second clinical evaluation at the age of 30 months or older to confirm their initial diagnosis (Table 2). Thus, all of the analyses reported herein were performed with participants whose autism (or typically developing) status was longitudinally tracked and confirmed at final diagnosis ages of 30 months or older.

**Table 1:** Subject characteristics table. Mean (standard deviation).

| <u>Characteristics</u> | <u>Autism</u><br>n= 68 |  | <u>Typically Developing Controls</u><br>n= 29 |
| --- | --- | --- | --- |
| Gender | 53 male, 15 female |  | 15 male, 14 female |
| Age at scan (months) | 31.5 (9) |  | 29.2 (10.2) |
|  | In proximity to scan | Final visit | In proximity to scan |
| ADOS Social & Communication Total * | 14.32 (3.9) | 14.4 (3.3) | 1.4 (1.4) |
| Vineland Communication | 76.76 (14.46) | 80 (17.8) | 105.4 (11.2) |
| Vineland Socialization | 79.62 (11.95) | 78.6 (14.5) | 108.2 (12.1) |
| Mullen Receptive Language T score | 27.88 (12.93) | 28.89 (13.5) | 56.3 (7.4) |
| Mullen Expressive Language T score | 29.11 (12.1) | 27.2 (17) | 56.7 (7.9) |
| * Participants received either the Toddler Module or Module 1 of the ADOS depending on their age and language ability at time of testing. |  |  |  |

**Table 2:** Initial and final ADOS Social and Communications scores of toddlers who were scanned at the age of 26 months old or younger. Two subjects, #4 and #15, were not identified as having autism at their early-age evaluations but received a final diagnosis of autism at ages 32 and 34 months respectively.

| Subject number | Age at scan (months) | ADOS assessment (in proximity to scan) |  | Final ADOS assessment |  |
| --- | --- | --- | --- | --- | --- |
|  |  | Age (months) | ADOS S&C total | Age (months) | ADOS S&C total |
| 1 | 13 | 13 | 14 | 37 | 8 |
| 2 | 14 | 13 | 15 | 34 | 14 |
| 3 | 15 | 14 | 17 | 45 | 18 |
| 4 | 15 | 14 | 3 | 32 | 16 |
| 5 | 16 | 14 | 5 | 39 | 19 |
| 6 | 18 | 16 | 18 | 33 | 10 |
| 7 | 20 | 19 | 13 | 42 | 19 |
| 8 | 20 | 19 | 19 | 35 | 15 |
| 9 | 20 | 20 | 14 | 32 | 18 |
| 10 | 20 | 20 | 20 | 38 | 11 |
| 11 | 21 | 20 | 18 | 34 | 16 |
| 12 | 21 | 21 | 15 | 44 | 11 |
| 13 | 23 | 22 | 18 | 34 | 13 |
| 14 | 24 | 24 | 18 | 45 | 14 |
| 15 | 24 | 24 | 0 | 34 | 18 |
| 16 | 25 | 24 | 12 | 47 | 13 |
| 17 | 25 | 24 | 12 | 34 | 8 |
| 18 | 26 | 26 | 15 | 40 | 19 |

### MRI acquisition

Anatomical T1 weighted and diffusion MRI scans were performed using a GE 1.5T Signa EXCITE scanner located at the UC San Diego Radiology Imaging Laboratory in Sorrento Valley, California. The scanner was equipped with an eight-channel phased-array head coil. The T1 weighted anatomical scan was acquired using a 3D fast spoiled gradient echo (FSPGR) sequence (166 sagittal slices: 0.94 * 0.94 * 1.2 mm, TE = 2.8ms, TR = 6.5ms, flip angle 12°, FOV = 24cm). Diffusion MRI was collected using a single-shot, echo-planar diffusion-weighted sequence (TE = 80.4ms, TR = 14500ms, flip angle 90°), which was applied along 51 diffusion directions. Data were collected from fifty 2.5mm thick axial slices (±10 slices, depending on head size; no gap) at in-plane resolution of 1.875 × 1.875 mm, using a b-value of 1000 sec/mm^2^. One non-diffusion weighted volume (b value=0) was acquired at the beginning of the scan for reference.

### Data Preprocessing

Data was preprocessed using mrDiffusion, an open source package written by the Vision, Imaging Science and Technology Activities (VISTA) lab at Stanford, CA, USA (http://web.stanford.edu/group/vista/cgi-bin/wiki/index.php/Software), and custom code written in Matlab (Mathworks, Natick, MA). The anterior commissure (AC), posterior commissure (PC), and mid-sagittal plane were identified manually and used to rotate the T1-weighted images to AC–PC aligned space via a rigid transform. Diffusion MRI data were corrected for head motion and eddy current distortions and registered to the non-diffusion weighted image (b0) using a nonlinear constrained transformation (Rohde et al. 2004). The b0 volume was registered to the T1 anatomy, using mutual information maximization methods implemented in SPM8 tools, and the same transformation was applied to all the diffusion weighted images. The gradient orientation matrix was transformed to compensate for all corrections and alignments in order to preserve the correct diffusion orientations (Leemans and Jones 2009). Finally, a tensor model was fit to the data in each voxel using a robust least-squares algorithm that removed outliers during tensor estimation (Robust Estimation of Tensors by Outlier Rejection, RESTORE (Chang et al. 2005)), and the 3 diffusion eigenvectors and eigenvalues of the tensor were extracted. Based on these measures, fractional anisotropy (FA), mean diffusivity (MD), radial diffusivity (RD), and axial diffusivity (AD) measures were calculated in each voxel (Basser and Pierpaoli 1996). AD is the first eigenvalue of the tensor, RD is the mean of the second and third eigenvalues, MD is the mean of all three eigenvalues, and FA is the normalized standard deviation of the three eigenvalues indicating the degree to which the tensor is anisotropic.

### Identification of the CC fiber groups

The Automated Fiber Quantification (AFQ) toolbox (Yeatman et al. 2012) was used to automatically identify seven CC tracts in each of the subjects. This procedure involved three steps: (1) whole brain tractography (2) region-of-interest (ROI) based segmentation, and (3) automated fiber tract cleaning.

1. Whole brain tractography was performed using a deterministic streamlines tracking algorithm (Mori et al. 1999; Basser et al. 2000). The tracking algorithm was seeded with a white matter mask of all voxels with FA values greater than 0.2. Starting from 8 seed points within each white matter voxel, path tracing proceeded in both directions in 1mm steps along the principal diffusion direction (PDD) of the tensor (i.e., the eigenvector with the largest eigenvalue). Tracking was stopped if the FA value dropped below 0.15, or if the angle between the diffusion direction in the current tensor and the direction in the next tensor was greater than 30°. This produced a large set of fibers throughout the entire brain for each toddler.
2. Fibers from step 1 that connect specific regions (e.g., the occipital lobes) of the two hemispheres through the CC were identified using pairs of homologous ROIs that were located in the white matter of the right and left hemispheres and a CC ROI in the mid sagittal plane, similarly to previous studies (Huang et al. 2005; Dougherty et al. 2007; Blecher et al. 2016). The seven pairs of symmetrical ROIs were defined on the Montreal Neurological Institute (MNI) template (Fonov et al. 2011) as follows:

a. Occipital tract – Two vertical ROIs included all white matter voxels located in the occipital lobe of coronal MNI slice #71 (Figure 1J, green).
b. Temporal tract – Two vertical ROIs included all white matter voxels located in the temporal lobe of coronal MNI slice #98 (Figure 1I, purple).
c. Posterior parietal tract – Two vertical ROIs included all white matter voxels located in the parietal lobe of coronal MNI slice #71 (Figure 1J, yellow).
d. Superior parietal tract – Two horizontal ROIs included all white matter voxels located in the parietal lobe of axial MNI slice #121 (Figure 1H, dark blue).
e. Posterior frontal tract – Two horizontal ROIs included white matter voxels located below the pre-central gyrus in the frontal lobe of axial MNI slice #121 (Figure 1H, pink).
f. Middle frontal – Two horizontal ROIs included all white matter voxels that were anterior to the pre-central gyrus in the frontal lobe of axial MNI slice #121 (Figure 1H, red).
g. Anterior frontal – Two vertical ROIs included all white matter voxels in the frontal lobe of coronal MNI slice #176 (Figure 1G, orange). Scans from each toddler were registered to the MNI template by computing a non-linear transformation between the b=0 diffusion volume of each toddler and the template, using a mutual information maximization algorithm (Friston 2004). We then applied the inverse transform to import the ROIs from template space into the native space of each toddler. Fibers from the whole brain tractography (step 1) that intersected the three relevant ROIs were labeled in each toddler (Figure 1). We were unable to identify fibers that intersected with the three relevant ROIs of the anterior frontal tract in two toddlers with autism, the posterior frontal tract in two other toddlers with autism, and the posterior parietal tract in one toddler with autism (5 different toddlers). All tracts were identified in all other toddlers.
3. Automated fiber tract cleaning was performed using an iterative procedure, which removed fibers that were more than 4 standard deviations above the mean fiber length as well as fibers located more than 5 standard deviations away from the center of the tract (see Yeatman et al., 2012).

**Figure 1.**
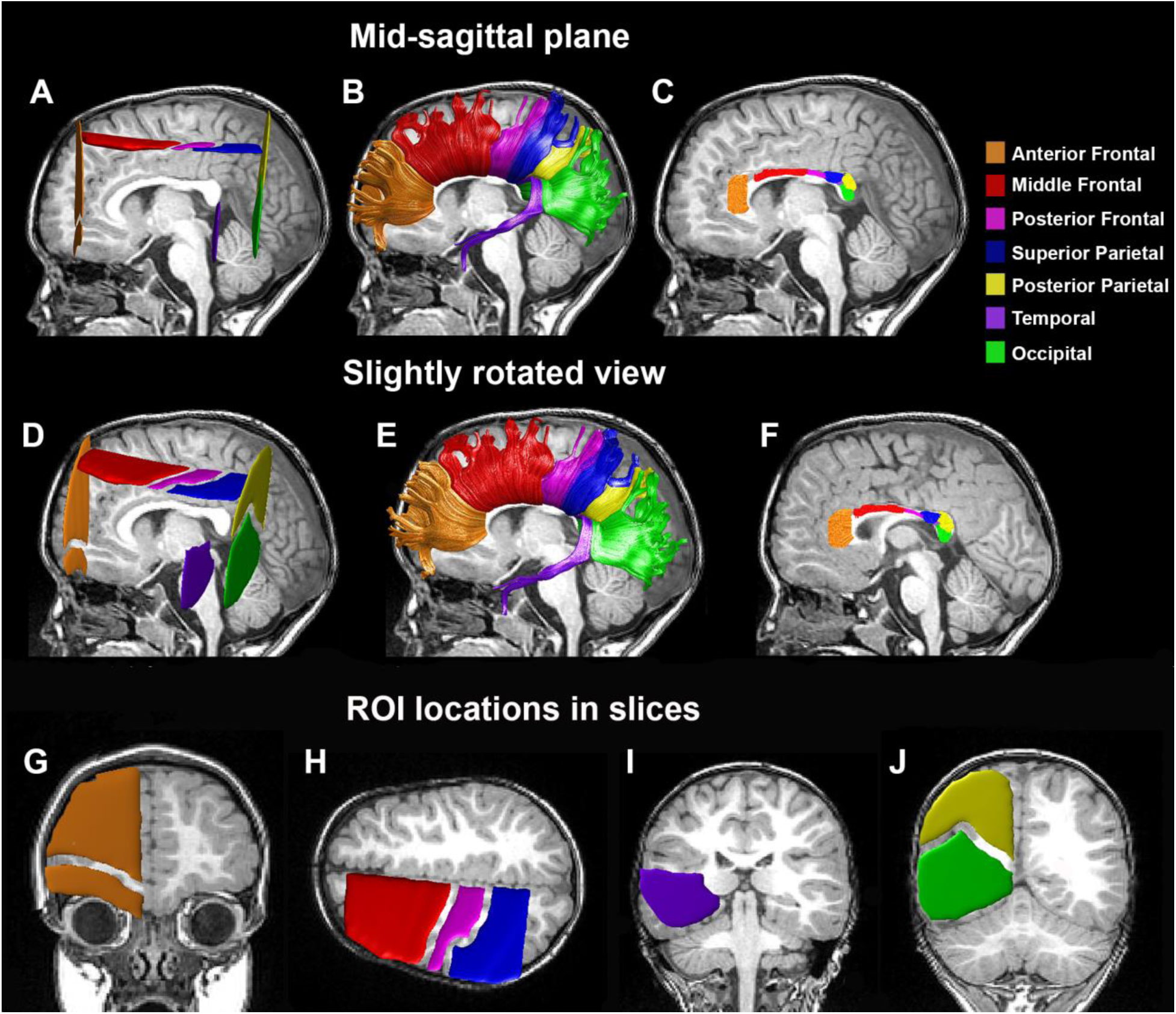
Fiber tract identification. (A,D) Left ROIs in an individual subject (male, 43 months old, with autism). Homologous ROIs were also defined in the right hemisphere. Tracts were identified by selecting fibers that intersected the left ROI, corpus callosum ROI, and homologous right ROI. (B,E) The seven CC fiber tracts that were identified in this individual. (C,F) Clipped segments of the CC fiber tracts. The same individual is presented in a mid-sagittal view (A-C) and a slightly rotated view (D-F) demonstrating the relative ROI locations in the left hemisphere. Diffusivity measures were extracted from the 10mm mid-sagittal segments presented in C and F. In addition, we present the specific coronal slice containing the anterior frontal ROI (G), the axial slice containing the middle-frontal, posterior frontal, and superior parietal ROIs (H), the coronal slice containing the temporal ROI (I), and the coronal slice containing the posterior parietal and occipital ROIs (J).

In a final step, we clipped the CC fiber tracts 5 mm to the right and to the left of the midline in order to focus the analyses on the diffusion properties in the medial section of the corpus callosum (Figure 1C), where directional coherence is maximal and there are no crossing or kissing fibers (Jeurissen et al. 2013). Rogue fibers that crossed the midsagittal plane outside a pre-defined bounding box covering each CC segment were excluded from the tracts before final analysis.

### Quantification of diffusion properties and cross-sectional area

The diffusion properties of each inter-hemispheric tract were calculated for each subject by computing the mean FA, MD, RD, and AD values across all voxels located in the 10mm midsagittal segment of that tract (corresponding to +/− 5mm from the midline). The cross sectional area of each callosal segment was computed by counting the number of anatomical voxels (1x1x1mm) that were traversed by the clipped fibers described above and dividing the result by 10mm (the segment length).

### Younger and older toddlers

Previous research has repeatedly shown that white-matter diffusion properties develop differently in toddlers with autism and controls such that significant differences are usually evident at earlier ages (Wolff et al. 2012; Solso et al. 2016). We therefore decided to perform a median split of the toddlers in both groups and created “younger” (Autism: n=32, Control n=16, age range: 13-30 months) and “older” (Autism: n=36, Control: n=13, age range: 31-45 months) age-matched subgroups. All measures were compared separately for each of the age groups.

### Relationship with behavioral measures

Each diffusion property was correlated with each of the behavioral measures to identify potential relationships with autism symptom severity (ADOS), adaptive social and communicative ability (Vineland), and language ability (Mullen). This analysis was performed only in the young autism group for the CC segments that exhibited significant differences across autism and control groups. The analysis was performed once using the behavioral measures that were collected within 12 weeks of the scan date and again using the measures acquired during the follow-up visit when autism diagnosis was confirmed (at the age of 2.5 years old or later). To ensure that potential findings were not due to the existence of outliers, we also examined these relationships using robust regression as implemented in Matlab. In this iterative analysis, a weighting vector is built, which discounts the effects of data points according to their leverage on the regression solution (i.e., by determining how much the regression solution changes before/after removing each point). Data points with extreme leverage (i.e. outliers) are thus ignored (Holland and Welsch 1977).

### Head motion

We estimated head motion in each subject using Framewise Displacement (FD) algorithm, which measures changes in head position from one volume to the next (e.g., Power et al., 2012). We computed the mean absolute translation and rotation FD across all volumes for each subject and then compared the results across the four groups. Furthermore, we performed a multiple regression analysis with the FD measures and each of the diffusion measures, extracted the residual diffusion values (after regressing out the FD measures), and re-ran the main analysis across younger autism and control toddlers. This analysis removed any potential between-subject differences in diffusion measures that were due to head motion differences across subjects.

### White matter volume

The white matter volume of each subject was estimated by counting the voxels with FA values greater than 0.2 in the entire brain.

### Statistical tests

Tract diffusion, cross-sectional area, head motion, white matter volume, and behavioral measures were compared across groups using two-tailed t-tests for independent measures with unequal variance. We used the false discovery rate (FDR) technique (Benjamini and Hochberg 1995) to correct the significance levels for multiple comparisons when examining measures per tract. Since MD and FA measures are based on AD and RD values, the statistical tests performed with MD and FA measures are not independent of those performed with the AD and RD values. We, therefore performed correction for multiple comparisons separately for AD/RD values and MD/FA values. The correction was performed across 14 comparisons – the two relevant diffusion measures (AD/RD or MD/FA) in each of the seven CC segments. The statistical significance of the Pearson's correlation coefficients, which were used to examine the relationship between the diffusion measures and the behavioral scores, were also corrected in the same manner. In this case FDR correction was performed across 4 comparisons – the two relevant diffusion measures (AD/RD or MD/FA) in each of the two examined CC segments that exhibited significant between-group differences (i.e., temporal and occipital tracts).

## Results

We compared the diffusion properties of seven callosal tracts between 68 toddlers with autism and 29 controls. We performed the analyses on the medial section of each tract encompassing 5 mm on either side of the mid-sagittal plane. This restricted the analyses to the segment with the maximal directional coherence (i.e. the most robust segment of each tract). To account for potential developmental changes across autism and control groups, we performed the analyses separately for toddlers younger or older than 2.5 years of age (see Methods).

In comparison to the young control toddlers, young toddlers with autism exhibited significantly larger FA values in the temporal CC segment (t(46) = 2.74, p =0.04, FDR corrected) and smaller MD, AD and RD values in the occipital and temporal segments (t(46) < −2.8, p < 0.04, FDR corrected, Figure 2 A-D). Note that similar differences were also apparent in the mid-frontal and anterior-frontal tracts, yet these differences did not survive correction for multiple comparisons. We did not find any significant between-group differences in any of the examined tracts in the older toddler subgroup (Figure 2 E-H).

**Figure 2.**
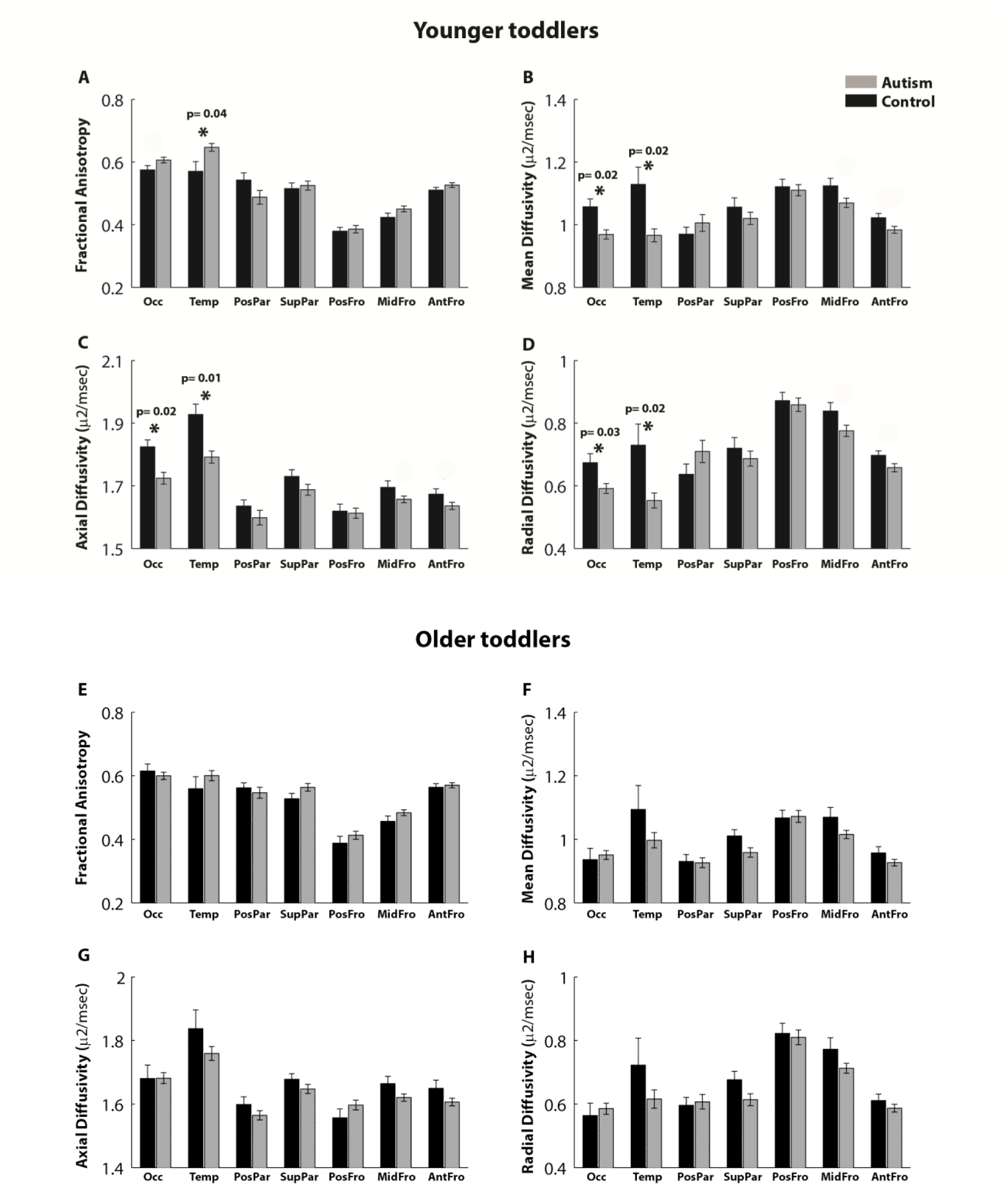
Comparison of diffusion properties in the seven CC tracts across the younger autism and control toddlers (<2.5 years old, A-D) and the older autism and control toddlers (>2.5 years old, E-H). Mean fractional anisotropy (A,E), Mean diffusivity (B,F), Axial diffusivity (C,G), and Radial diffusivity (D,H) are presented for the autism (gray) and control (black) groups. Error bars: standard error of the mean across subjects. Asterisks: significant differences across groups after FDR correction (specific p values are noted for significant tracts).

Equivalent results were found when including only young male toddlers in the analysis, demonstrating that the results were not driven by gender differences across groups. Young male toddlers with autism exhibited significantly larger FA values in the temporal CC segment (t(29)= 2.8, p= 0.04, FDR corrected), smaller MD and AD values in the occipital and temporal segments (t(29)< −3, p< 0.04, FDR corrected), and smaller RD values in the temporal segment (t(29)=-3.1, p=0.02, FDR corrected) in comparison to young male control toddlers.

An examination of the cross-sectional area of each tract (see Methods) also revealed significant between-group differences only in the younger toddler subgroup, where the temporal tract was larger in toddlers with autism (t(46) =3.12, p =0.02, FDR corrected, Figure 3, left). There were no significant between-group differences in any of the tracts in the older subgroup (Figure 3, right).

**Figure 3.**
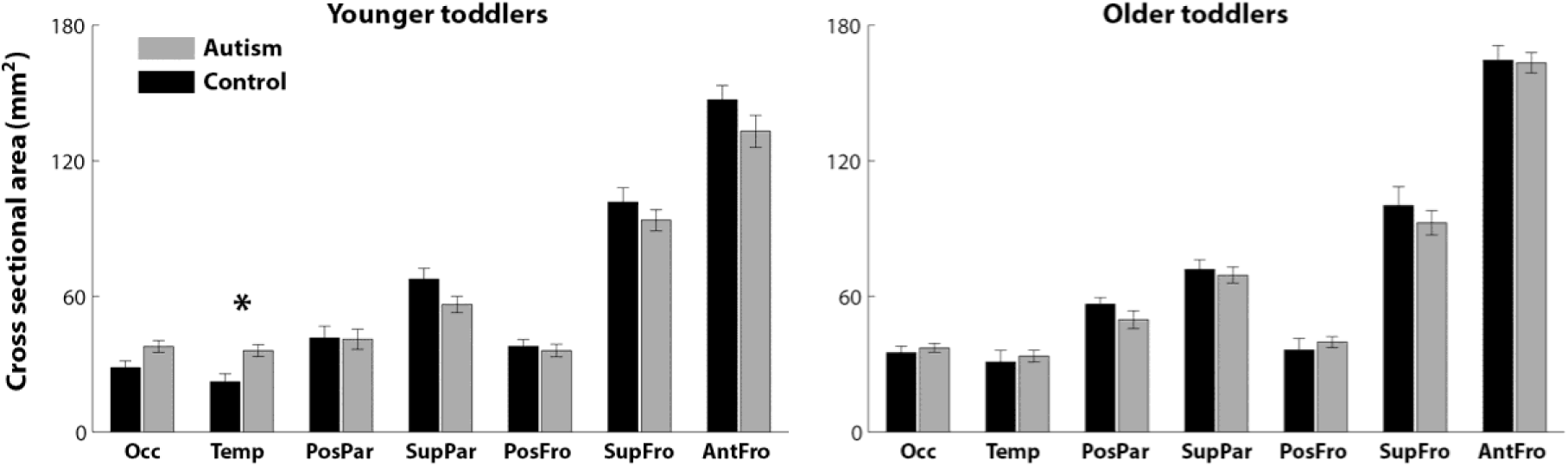
Comparison of tract cross-sectional area in the younger autism and control toddlers (left panel) and in the older autism and control toddlers (right panel). Gray: Autism, Black: Control. Error bars: standard error of the mean across subjects. Asterisks: significant differences after FDR correction (p < 0.05).

Correlations between diffusion measures and behavioral measures were calculated only for the younger toddlers with autism in the tracts that exhibited significant between-group differences in diffusion measures (i.e., the occipital and temporal tracts, Figure 4). Pearson's correlations between each of the diffusion measures and the early behavioral scores measured that were acquired close to the scan date were not significant in any of the examined tracts (r (30) < 0.3, p > 0.05, uncorrected to increase sensitivity). However, Vineland-socialization scores acquired at the final outcome visit (6-30 months after the scan) exhibited significant negative correlations with FA values (r (30) = −0.5, p =0.04, FDR corrected) and significant positive correlations with MD (r (30) = 0.41, p = 0.05, FDR corrected) and RD values (r (30) = 0.46, p = 0.04, FDR corrected) in the temporal tract. Pearson's correlations between FA values and the ADOS scores were marginally significant for the temporal tract (r (30) = 0.42, p = 0.08, FDR corrected). Diffusion measures in the occipital tract were not correlated with behavioral measures. Note that ADOS scores increase with symptom severity while Vineland scores increase with social proficiency. The correlations between the cross sectional area of the temporal tract and the ADOS (r (30) =0.18, p =0.31, uncorrected) or Vineland (r (30) =-0.05, p =0.77, uncorrected) scores were not significant.

**Figure 4.**
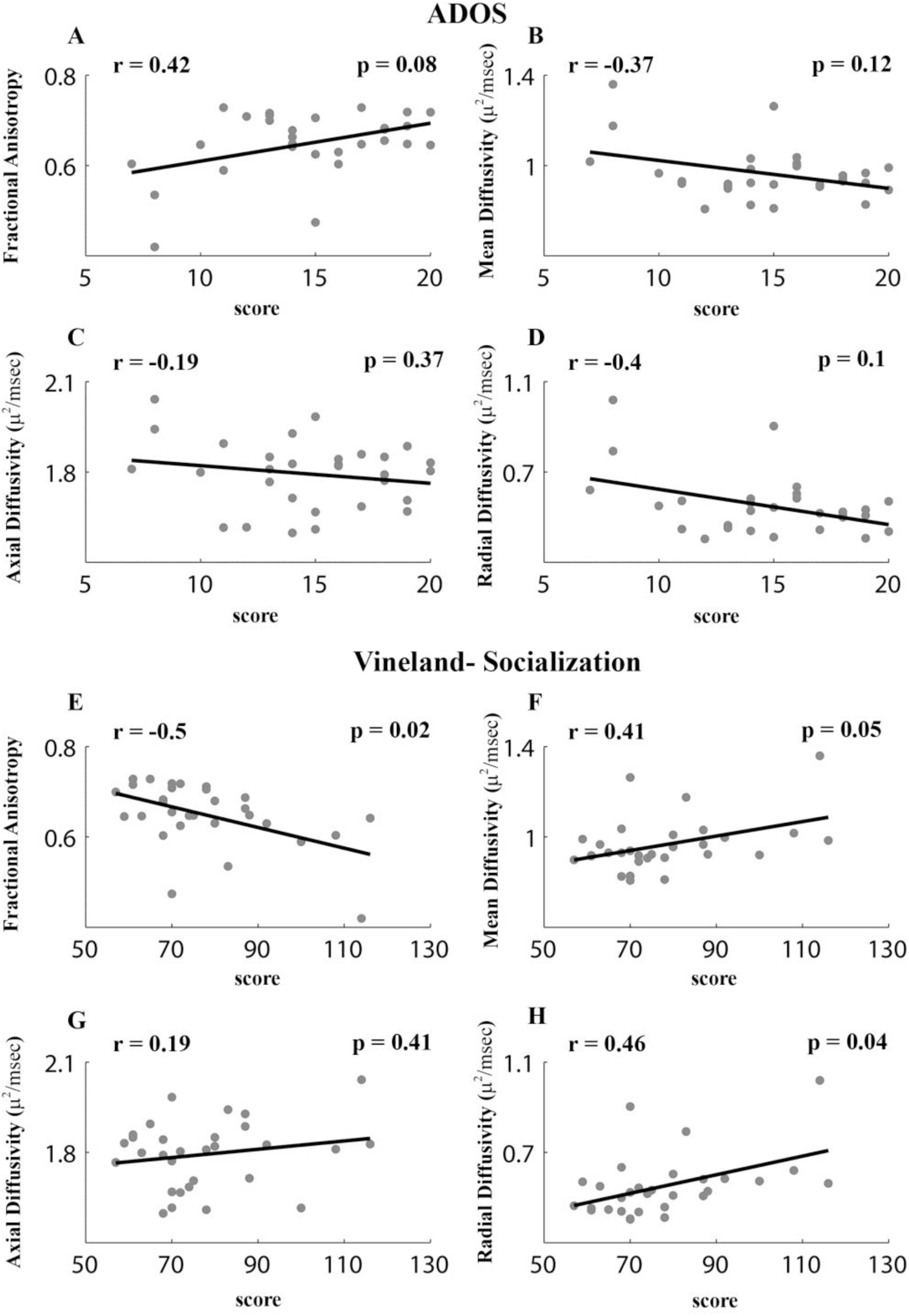
Scatter plots presenting the relationship between ADOS social and communication total scores (**A-D**) or Vineland socialization scores (**E-H**) at final diagnosis age and each of the diffusion measures in the temporal tract: Fractional anisotropy (A,E), Mean diffusivity (B,F), Axial diffusivity (C,G), and Radial diffusivity (D,H). Each point represents a single toddler. Only younger toddlers with autism (<2.5 years old, N=34) were included in this analysis.

To further examine the robustness of these relationships we also performed analogous robust regression analyses with diffusion measures in the temporal tract. These analyses yielded similar results when examining the Vineland-socialization scores, which exhibited significant negative relationships with FA values (β = −0.002, t(30) = −2.88, p = 0.007) and positive relationships with RD values (β = 0.002, t(30) = 2.5, p = 0.02). Analyses with ADOS scores did not yield significant results (p > 0.1). This indicates that the relationship between diffusion measures and Vineland scores was robust to the existence of potential outliers in the data, but the relationship with ADOS scores was not (see Methods).

Pearson's correlation coefficients and their significance are noted in each panel. All p-values are FDR corrected.

We performed several control analyses to rule out alternative interpretations of the results described above. First, we examined estimated head motion parameters using the mean framewise displacement (FD) measure (see Methods), which was computed separately for each of the six head rotation and translation estimates in each subject. Younger toddlers from both the autism and control groups exhibited larger FD values than older toddlers of the corresponding group (t(66) and t(27) > 2, p < 0.05), but there were no significant differences between the autism and control toddlers within each age group (t(46) and t(47) < 1, p > 0.3, Figure 5A). The same was true when comparing the entire sample (t(95) > −0.5, p > 0.6). This means that toddlers with autism did not move their heads significantly more than controls. In addition, we also re-ran our main analysis after first regressing-out the estimated head-motion parameters of each subject from the diffusion measures (see Methods) and found similar results. Younger toddlers with autism exhibited significantly larger FA values in the temporal CC segment (t(46) = 2.4, p = 0.03) and smaller MD, AD, and RD values in the occipital and temporal CC segments (t(46)< −2.1, p< 0.05). These analyses rule out the possibility that differences in diffusion measures across groups were caused by differences in head motion during MRI scanning.

**Figure 5.**
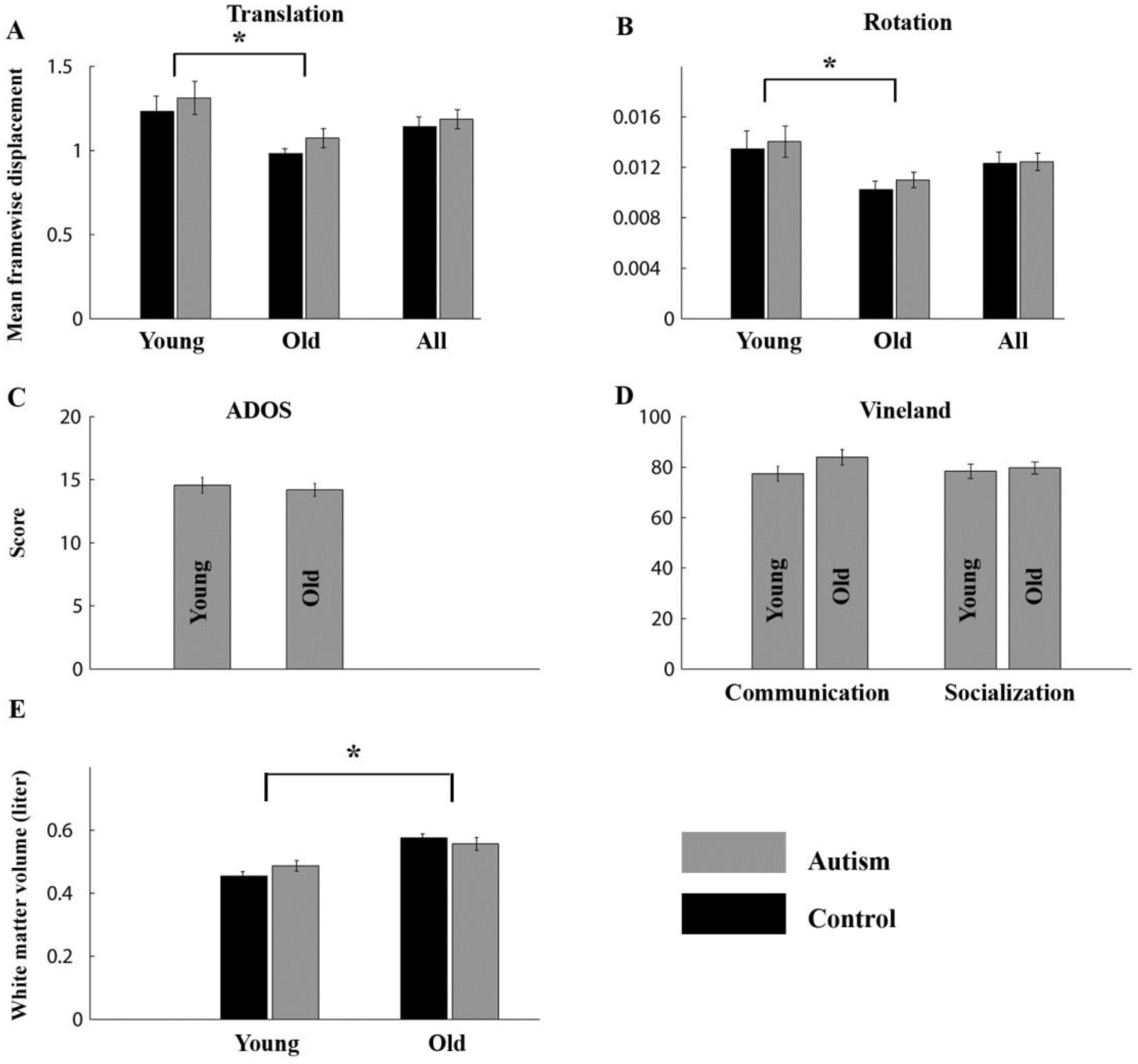
Control analyses. Mean framewise displacement of head rotations (A) and translations (B) were compared across groups when including all toddlers (All) and after splitting younger and older toddlers. (C) Comparison of ADOS social and communication total scores across younger and older toddlers with autism. (D) Comparison of Vineland communication and socialization scores across younger and older toddlers with autism. (E) Comparison of overall white matter volume across autism and control toddlers. Gray: Autism, Black: Control. Error bars: standard error of the mean across subjects. Asterisks: significant differences (p < 0.05, uncorrected to increase sensitivity).

Second, we examined whether there were differences in the severity of autism symptoms across age groups and found no significant differences in either ADOS or Vineland scores between the younger and older toddlers with autism (t(66) = 0.5 and t(66) = 0.1 respectively, p > 0.1, Figure 5B). This ensured that the specificity of findings to the young toddler groups was not generated by differences in autism severity between younger and older toddlers with autism.

Third, we examined overall white matter volumes in each of the groups (see methods). As expected, younger toddlers from both autism and control groups had significantly smaller white matter volumes than older toddlers from each group (t(66) and t(27) > 4, p < 0.0005), but there were no significant differences across autism and control toddlers within each age group (t(46) and t(47) < 2, p > 0.2, Figure 5C). This analysis rules out the possibility that differences in diffusion measures across groups were generated by differences in overall white matter volumes across autism and control toddlers.

## Discussion

At the earliest age of clinical onset, 1 to 2.5 year old toddlers with autism already exhibit multiple structural abnormalities in callosal tracts connecting the temporal lobes as compared with matched typically developing toddlers (Figure 2). Larger FA and smaller MD, AD, and RD values were clearly apparent in this tract, which was significantly larger in cross-sectional area (Figure 3). Smaller diffusion values (MD, AD, and RD) and larger directionality (FA) suggest that the temporal tract may contain an abnormally large number of small caliber, densely packed axons (see below) that span a larger cross-sectional area of the CC. Moreover, the degree of micro-structural abnormality in the temporal callosal pathway at the age of clinical ***<u>onset</u>*** was correlated with the severity of social clinical symptoms as measured by the Vineland test at ***<u>later outcome ages</u>*** (marginally significant also with ADOS scores, Figure 4). This suggests that early anatomical abnormalities may be useful for assessing clinical outcome and prognosis. Finally, the micro-structural abnormalities apparent in the younger toddlers with autism (<2.5 years old) were not evident in the older toddlers with autism (>2.5 years old). Nevertheless, we speculate that dynamic, ongoing pathology manifested in the number, size, density, and/or myelination of CC axons disrupts typical inter-hemispheric connectivity across the temporal lobes and may be a common characteristic of many individuals with autism not only during early development, but also later in life.

### Nature of micro-structural CC abnormalities in autism

Larger FA values along with smaller MD, AD, and RD values indicate that water diffusion is more restricted in the temporal CC tracts of young toddlers with autism as compared with the young control toddlers. Several differences in white matter micro-structure may yield this combination of diffusion values, including increased myelination, increased axonal density, smaller axonal calibers, and/or increased extra-cellular matrix density (Mori and Zhang 2006). Numerous postmortem (Courchesne and Pierce 2005; Courchesne, Mouton, et al. 2011; Santos et al. 2011; Stoner et al. 2014), genetic (Pinto et al. 2010; Chow et al. 2012; Bernier et al. 2014; Sugathan et al. 2014; Pramparo et al. 2015), animal model (Fang et al. 2014; Le Belle et al. 2014; Orosco et al. 2014; Sabers et al. 2014), and cellular (Marchetto et al. 2016; Packer 2016) studies suggest that, in many cases, autism is associated with a prenatal excess of neural proliferation, which would predict an excess of corresponding axons. While postmortem studies with toddlers have not examined axonal characteristics or reported axon numbers in the white matter of toddlers with autism, adults with autism do exhibit an abnormally large number of smaller caliber axons in frontal lobe areas (Zikopoulos and Barbas 2010). One plausible explanation for our DTI findings, therefore, is that they reflect an underlying physiology of excessive neuron numbers in temporal areas, which send out a corresponding large number of small caliber callosal axons. Such developmental alterations are likely to be of prenatal origin and reflect abnormalities in cell proliferation, migration, differentiation and axon and dendritic development (Courchesne, Mouton, et al. 2011; Chow et al. 2012; Stoner et al. 2014).

### Timing of micro-structural CC abnormalities in autism

Abnormally high FA and low MD, AD, and RD values seem to appear transiently at the very early stages of autism development when clinical symptoms initially emerge. In our sample, only the younger toddler group (<2.5 years old) exhibited significant diffusivity differences in comparison to their matched control group while the older toddler group (<2.5 years old) did not. This conclusion is supported by several other DTI studies that have reported significantly larger FA values in callosal and/or other white matter tracts in young toddlers with autism (Ben Bashat et al. 2007; Weinstein et al. 2011; Wolff et al. 2013). The studies that examined the effects of age reported that the abnormally high FA values found in toddlers who develop autism at young ages (0.5-3 years old) were transient and gradually disappeared or even reversed to abnormally low FA values during early development (Ben Bashat et al. 2007; Weinstein et al. 2011; Travers et al. 2015). One study that compared high risk-sibs who eventually developed autism to those who did not, reported that the transition from abnormally high to low FA values in the autism group took place before toddlers reached the age of 1.5 years old (Wolff et al. 2012). Two other studies that compared toddlers with autism to control toddlers reported that this transition happens a little later, around the age of three (Weinstein et al. 2011; Solso et al. 2016). Our results are in line with the latter studies and suggest that significant findings in the CC are only apparent in toddlers who are younger than 2.5 years old. Taken together, it seems that the choice of comparison group (typically developing versus high risk sibs who may share some of the etiological factors exhibited by toddlers with autism) affects the reported developmental changes in FA values.

In agreement with these findings, DTI studies with older children (Nordahl et al. 2015; Travers et al. 2015; Wolff et al. 2015), adolescents (Alexander et al. 2007), and adults (Catani et al. 2008, 2016) have commonly reported either non-significant differences across groups, differences in the variability of FA values within the brain (Cascio et al. 2013), or abnormally low FA values in individuals with autism. This highlights the importance of studying autism neurobiology during early development (Courchesne et al. 2007; Pierce et al. 2011).

While abnormal diffusivity measures appear transiently during early autism development, inter-hemispheric connectivity abnormalities may persist throughout life as suggested by fMRI studies that have reported poor functional synchronization across the hemispheres (Anderson et al. 2011; Di Martino et al. 2014) in adolescents and adults with autism. When attempting to reconcile different types of evidence, it is important to keep in mind that the ability to measure inter-hemispheric connectivity with specific tools may change as a consequence of the dramatic multi-factorial changes that take place during early development.

### Location of micro-structural CC abnormalities in autism

One advance of the current study was the use of deterministic tractography to sub-divide the CC into seven fiber groups (i.e., inter-hemispheric tracts) according to the projections of the fibers in specific cortical areas. Separating CC tracts in this manner revealed that abnormal micro-structure during early autism development was localized to temporal and occipital tracts. Moreover, abnormalities in the temporal tract were apparent also in the cross-sectional area of the tract and diffusion abnormalities were correlated with the Vinland socialization measures (and marginally correlated with the ADOS scores, Figure 4), suggesting that the development of this tract is of particular interest.

Previous DTI studies that examined CC micro-structure in young toddlers with autism also applied deterministic tractography techniques (Nordahl et al. 2015; Travers et al. 2015; Wolff et al. 2015), but used relatively crude methods to divide the CC into three segments without examining the cortical projections of the CC fibers. In these studies, the examined tracts were identified using ROIs located in the genu, body, and/or splenium of the CC without the use of additional white matter ROIs that delineate tracts based on their projections into specific cortical regions. This means that study-specific demarcations of the three CC areas were likely to include different combinations of inter-hemispheric tracts that connect multiple cortical regions. With this in mind, significantly increased FA in toddlers with autism was reported only in the body of the CC in one of the studies (Wolff et al. 2012), only in the genu in another study (Solso et al. 2016), and in both CC segments in the third (Travers et al. 2015). Our results present a more detailed localization of the affected inter-hemispheric tracts, which, in contrast to previous studies, highlights the role of temporal CC tract abnormalities during early autism development. Additional studies utilizing cortical ROIs and probabilistic tractography techniques will be useful for further examining the specificity of affected tracts and the robustness of the results across multiple tractography techniques.

### Potential relationship with functional connectivity

Micro-structural CC abnormalities are likely to be associated with poor inter-hemispheric functional synchronization. An overabundance of small caliber CC axons is likely to alter neural communication across the hemispheres in complex ways. For example, smaller caliber axons will lead to slower neural transmission speeds (Laughlin and Sejnowski 2003), which may impair the ability of bi-lateral neural populations to synchronize their activity. Several autism studies with adolescents, adults (Anderson et al. 2011; Di Martino et al. 2014), and toddlers (Dinstein et al. 2011) have indeed reported that individuals with autism exhibit poor inter-hemispheric synchronization (weak functional connectivity). In naturally sleeping toddlers, poor inter-hemispheric synchronization seems to be most apparent between right and left superior temporal gyri (the general location of Wernike's area) (Dinstein et al. 2011). Taken together with the current results, we hypothesize that structural abnormalities in the temporal CC tract may be associated with impaired functional connectivity across temporal cortical areas during early autism development. Relating these functional and structural measures within the same toddlers will be critical for elucidating the underlying neuropathology of affected toddlers.

### Potential behavioral consequences

The significant correlations between Vineland-socialization scores and diffusivity measures in the temporal CC tract, suggest that micro-structural abnormalities in this tract may be of particular behavioral importance. We speculate that impaired inter-hemispheric connectivity across the temporal lobes may have specific consequences for the development of lateralized functions such as language and social communication. Several studies have reported that temporal lobe responses to language stimuli are abnormally lateralized in toddlers with autism (Redcay and Courchesne 2008; Eyler et al. 2012) and that toddlers with autism who exhibit weak temporal cortex responses to language stimuli early on, develop poor language capabilities later in life, while those with near-typical responses develop good language capabilities (Lombardo et al. 2015). Similarly, teenagers who had been born preterm and developed language impairments exhibit smaller interhemispheric white matter fibers across temporal lobes in comparison to those who do not develop language impairments (Northam et al. 2012). Adults who had been born preterm exhibit lower FA values in posterior segments of the CC that are associated with abnormal fMRI responses during verbal learning tasks in comparison to controls (Salvan et al. 2014) .

Studies with typically developing adults have suggested that specific, specialized areas in temporal cortex including the superior temporal sulcus (STS) (Allison et al. 2000; Zilbovicius et al. 2006) and temporo-parietal junction (TPJ) (Saxe 2006) are critical for social interactions. Adolescents and adults with autism indeed exhibit abnormally weak fMRI responses in the STS during processing of eye gaze (Pelphrey et al. 2005) and biological motion (Herrington et al. 2007; Freitag et al. 2008) and abnormally lateralized TPJ responses during judgments of mental state (Lombardo et al. 2011). Relating abnormalities in structural and functional inter-hemispheric connectivity with abnormalities in functional specialization of these cortical areas (within the same individuals) seems like another important goal for future autism research.

### Heterogeneity in autism

It is important to note that individuals with autism differ from one another in terms of their behavioral profiles, intelligence, clinical comorbidities, genetics, brain structure, and brain function. This variability across participants was clearly apparent in the current study (Figure 4). When interpreting our results, we believe that it is, therefore, more useful to consider inter-hemispheric connectivity abnormalities as an indicator of one potential pathological mechanism that may be present in some, but not all, toddlers with autism. With this in mind it seems more promising to conduct future studies while separating the autism population into subgroups that exhibit common structural, functional (Lombardo et al. 2015), eye tracking (e.g., (Pierce et al. 2015), behavioral, genetic, sensory, and/or clinical findings.

### Conclusions

Multiple structural abnormalities were clearly apparent in the CC tracts that connect the temporal lobes of 1 to 2.5 year old toddlers who developed autism. We suggest that these toddlers may have an overabundance of small, tightly packed axons, which may impair inter-hemispheric communication and lead to poor functional synchronization and specialization. Additional research focusing on early inter-hemispheric connectivity while combining structural and functional MRI techniques within the same toddlers will be critical for elucidating the underlying neuropathology.

## Acknowledgments

This study was supported by NIMH grant P50-MH081755 (E.C.), NIMH grant R01-MH36840 (E.C.), NIMH grant R01-MH080134 (K.P.), Autism Speaks (K.P.), Cure Autism Now (K.P.), ISF grant 961/14 (I.D.), and GIF grant 2351 (I.D.). The authors thank S. Solso, K. Campbell, C. Ahrens-Barbeau, J.Young, M. Mayo, and S. Marinero for help with data collection, and all families who participated in research at the UC San Diego Autism Center. The authors have no conflicts of interest with regard to the presented work.

## References

Alexander AL, Lee JE, Lazar M, Boudos R, DuBray MB, Oakes TR, Miller JN, Lu J, Jeong EK, McMahon WM, Bigler ED, Lainhart JE. 2007. Diffusion tensor imaging of the corpus callosum in Autism. Neuroimage. 34:61–73.)

Allison T, Puce A, McCarthy G. 2000. Social perception from visual cues: Role of the STS region. Trends Cogn Sci. 4:267–278.

Ameis SH, Catani M. 2015. Altered white matter connectivity as a neural substrate for social impairment in Autism Spectrum Disorder. Cortex. 62:158–181.

Anderson JS, Druzgal TJ, Froehlich A, Dubray MB, Lange N, Alexander AL, Abildskov T, Nielsen J a., Cariello AN, Cooperrider JR, Bigler ED, Lainhart JE. 2011. Decreased interhemispheric functional connectivity in autism. Cereb Cortex. 21:1134–1146.

Assaf Y, Pasternak O. 2008. Diffusion tensor imaging (DTI)-based white matter mapping in brain research: a review. J Mol Neurosci. 34:51–61.

Barnea-Goraly N, Kwon H, Menon V, Eliez S, Lotspeich L, Reiss AL. 2004. White matter structure in autism: Preliminary evidence from diffusion tensor imaging. Biol Psychiatry. 55:323–326.

Basser PJ, Pajevic S, Pierpaoli C, Duda J, Aldroubi A. 2000. In vivo fiber tractography using DT-MRI data. Magn Reson Med. 44:625–632.

Basser PJ, Pierpaoli C. 1996. Microstructural and physiological features of tissues elucidated by quantitative-diffusion-tensor MRI. J Magn Reson B. 111:209–219.

Ben Bashat D, Kronfeld-Duenias V, Zachor D a., Ekstein PM, Hendler T, Tarrasch R, Even A, Levy Y, Ben Sira L. 2007. Accelerated maturation of white matter in young children with autism: A high b value DWI study. Neuroimage. 37:40–47.

Benjamini Y, Hochberg Y. 1995. Controlling the False Discovery Rate: A Practical and Powerful Approach to Multiple Testing. J R Stat Soc Ser B. 57:289–300.

Bernier R, Golzio C, Xiong B, Stessman HA, Coe BP, Penn O, Witherspoon K, Gerdts J, Baker C, Vulto-van Silfhout AT, Schuurs-Hoeijmakers JH, Fichera M, Bosco P, Buono S, Alberti A, Failla P, Peeters H, Steyaert J, Vissers LELM, Francescatto L, Mefford HC, Rosenfeld JA, Bakken T, O’Roak BJ, Pawlus M, Moon R, Shendure J, Amaral DG, Lein E, Rankin J, Romano C, de Vries BBA, Katsanis N, Eichler EE. 2014. Disruptive CHD8 Mutations Define a Subtype of Autism Early in Development. Cell. 158:263–276.

Blanchard DC, Defensor EB, Meyza KZ, Pobbe RLH, Pearson BL, Bolivar VJ, Blanchard RJ. 2012. BTBR T+tf/J mice: Autism-relevant behaviors and reduced fractone-associated heparan sulfate. Neurosci Biobehav Rev.

Blecher T, Tal I, Ben-Shachar M. 2016. White matter microstructural properties correlate with sensorimotor synchronization abilities. Neuroimage. 138:1–12.

Bolivar VJ, Walters SR, Phoenix JL. 2007. Assessing autism-like behavior in mice: Variations in social interactions among inbred strains. Behav Brain Res. 176:21–26.

Cascio C, Gribbin M, Gouttard S, Smith RG, Jomier M, Field S, Graves M, Hazlett HC, Muller K, Gerig G, Piven J. 2013. Fractional anisotropy distributions in 2- to 6-year-old children with autism. J Intellect Disabil Res. 57:1037–1049.

Catani M, Dell’Acqua F, Budisavljevic S, Howells H, Thiebaut de Schotten M, Froudist-Walsh S, D’Anna L, Thompson A, Sandrone S, Bullmore ET, Suckling J, Baron-Cohen S, Lombardo M V, Wheelwright SJ, Chakrabarti B, Lai M-C, Ruigrok AN V, Leemans A, Ecker C, Consortium MA, Craig MC, Murphy DGM. 2016. Frontal networks in adults with autism spectrum disorder. Brain. 139:616–630.

Catani M, Jones DK, Daly E, Embiricos N, Deeley Q, Pugliese L, Curran S, Robertson D, Murphy DGM. 2008. Altered cerebellar feedback projections in Asperger syndrome. Neuroimage. 41:1184–1191.

Chang L-C, Jones DK, Pierpaoli C. 2005. RESTORE: robust estimation of tensors by outlier rejection. Magn Reson Med. 53:1088–1095.

Chow ML, Pramparo T, Winn ME, Barnes CC, Li HR, Weiss L, Fan JB, Murray S, April C, Belinson H, Fu XD, Wynshaw-Boris A, Schork NJ, Courchesne E. 2012. Age-dependent brain gene expression and copy number anomalies in autism suggest distinct pathological processes at young versus mature ages. PLoS Genet. 8.

Courchesne E, Campbell K, Solso S. 2011. Brain growth across the life span in autism: Age-specific changes in anatomical pathology. Brain Res. 1380:138–145.

Courchesne E, Karns CM, Davis HR, Ziccardi R, Carper RA, Tigue ZD, Chisum HJ, Moses P, Pierce K, Lord C, Lincoln AJ, Pizzo S, Schreibman L, Haas RH, Akshoomoff NA, Courchesne RY. 2001. Unusual brain growth patterns in early life in patients with autistic disorder: an MRI study. Neurology. 57:245–254.

Courchesne E, Mouton PR, Calhoun ME, Semendeferi K, Ahrens-Barbeau C, Hallet MJ, Barnes CC, Pierce K. 2011. Neuron number and size in prefrontal cortex of children with autism. JAMA. 306:2001–2010.

Courchesne E, Pierce K. 2005. Why the frontal cortex in autism might be talking only to itself: local over-connectivity but long-distance disconnection. Curr Opin Neurobiol. 15:225–230.

Courchesne E, Pierce K, Schumann CM, Redcay E, Buckwalter J a, Kennedy DP, Morgan J. 2007. Mapping early brain development in autism. Neuron. 56:399–413.

Di Martino a, Yan C-G, Li Q, Denio E, Castellanos FX, Alaerts K, Anderson JS, Assaf M, Bookheimer SY, Dapretto M, Deen B, Delmonte S, Dinstein I, Ertl-Wagner B, Fair D a, Gallagher L, Kennedy DP, Keown CL, Keysers C, Lainhart JE, Lord C, Luna B, Menon V, Minshew NJ, Monk CS, Mueller S, Müller R, Nebel MB, Nigg JT, O’Hearn K, Pelphrey K a, Peltier SJ, Rudie JD, Sunaert S, Thioux M, Tyszka JM, Uddin LQ, Verhoeven JS, Wenderoth N, Wiggins JL, Mostofsky SH, Milham MP. 2014. The autism brain imaging data exchange: towards a large-scale evaluation of the intrinsic brain architecture in autism. Mol Psychiatry. 19:659–667.

Dinstein I, Pierce K, Eyler L, Solso S, Malach R, Behrmann M, Courchesne E. 2011. Disrupted Neural Synchronization in Toddlers with Autism. Neuron. 70:1218–1225.

Dougherty RF, Ben-Shachar M, Deutsch GK, Hernandez A, Fox GR, Wandell B a. 2007. Temporal-callosal pathway diffusivity predicts phonological skills in children. Proc Natl Acad Sci U S A. 104:8556–8561.

Egaas B, Courchesne E, Saitoh O. 1995. Reduced Size of Corpus Callosum in Autism. Arch Neurol. 52:794–801.

Eyler LT, Pierce K, Courchesne E. 2012. A failure of left temporal cortex to specialize for language is an early emerging and fundamental property of autism. Brain. 135:949–960.

Fang W-Q, Chen W-W, Jiang L, Liu K, Yung W-H, Fu AKY, Ip NY. 2014. Overproduction of Upper-Layer Neurons in the Neocortex Leads to Autism-like Features in Mice. Cell Rep. 9:1635–1643.

Fonov V, Evans AC, Botteron K, Almli CR, McKinstry RC, Collins DL. 2011. Unbiased average age-appropriate atlases for pediatric studies. Neuroimage. 54:313–327.

Frazier TW, Hardan AY. 2009. A meta-analysis of the corpus callosum in autism. Biol Psychiatry. 66:935–941.

Freitag CM, Konrad C, Häberlen M, Kleser C, von Gontard A, Reith W, Troje NF, Krick C. 2008. Perception of biological motion in autism spectrum disorders. Neuropsychologia. 46:1480–1494.

Friston. 2004. Generative and recognition models for neuroanatomy. Neuroimage. 23:17–20.

Haar S, Berman S, Behrmann M, Dinstein I. 2014. Anatomical Abnormalities in Autism? Cereb Cortex. 1–13.

Herrington JD, Baron-Cohen S, Wheelwright SJ, Singh KD, Bullmore ET, Brammer M, Williams SCR. 2007. The role of MT+/V5 during biological motion perception in Asperger Syndrome: An fMRI study. Res Autism Spectr Disord. 1:14–27.

Holland PW, Welsch RE. 1977. Robust regression using iteratively reweighted least-squares. Commun Stat - Theory Methods. 6:813–827.

Huang H, Zhang J, Jiang H, Wakana S, Poetscher L, Miller MI, Van Zijl PCM, Hillis AE, Wytik R, Mori S. 2005. DTI tractography based parcellation of white matter: Application to the mid-sagittal morphology of corpus callosum. Neuroimage. 26:195–205.

Jeurissen B, Leemans A, Tournier J-D, Jones DK, Sijbers J. 2013. Investigating the prevalence of complex fiber configurations in white matter tissue with diffusion magnetic resonance imaging. Hum Brain Mapp. 34:2747–2766.

Jou RJ, Mateljevic N, Kaiser MD, Sugrue DR, Volkmar FR, Pelphrey K a. 2011. Structural neural phenotype of autism: Preliminary evidence from a diffusion tensor imaging study using tract-based spatial statistics. Am J Neuroradiol. 32:1607–1613.

Just MA, Cherkassky VL, Keller T a., Minshew NJ. 2004. Cortical activation and synchronization during sentence comprehension in high-functioning autism: Evidence of underconnectivity. Brain. 127:1811–1821.

Laughlin SB, Sejnowski TJ. 2003. Communication in neuronal networks. Science. 301:1870–1874.

Le Belle JE, Sperry J, Ngo A, Ghochani Y, Laks DR, López-Aranda M, Silva AJ, Kornblum HI. 2014. Maternal Inflammation Contributes to Brain Overgrowth and Autism-Associated Behaviors through Altered Redox Signaling in Stem and Progenitor Cells. Stem Cell Reports. 3:725–734.

Leemans A, Jones DK. 2009. The B-matrix must be rotated when correcting for subject motion in DTI data. Magn Reson Med. 61:1336–1349.

Lombardo MV, Pierce K, Eyler LT, Carter Barnes C, Ahrens-Barbeau C, Solso S, Campbell K, Courchesne E. 2015. Different Functional Neural Substrates for Good and Poor Language Outcome in Autism. Neuron. 86:567–577.

Lombardo M V., Chakrabarti B, Bullmore ET, Baron-Cohen S. 2011. Specialization of right temporo-parietal junction for mentalizing and its relation to social impairments in autism. Neuroimage. 56:1832–1838.

Lord C, Risi S, Lambrecht L, Cook EHJ, Leventhal BL, DiLavore PC, Pickles a, Rutter M. 2000. The Autism Diagnostic Schedule – Generic: A standard measures of social and communication deficits associated with the spectrum of autism. J Autism Dev Disord. 30:205–223.

Marchetto M, Belinson H, Tian Y, Freitas B, Fu C, Vadodaria K, Beltrao-Braga P, Trujillo C, Mendes A, Padmanabhan K, Nunez Y, Ou J, Ghosh H, Wright R, Brennand K, Pierce K, Eichenfield L, Pramparo T, Eyler L, Barnes C, Courchesne E, Geschwind D, Gage F, Wynshaw-Boris A, Muotri A. 2016. Altered proliferation and networks in neural cells derived from idiopathic autistic individuals. Mol Psychiatry. In press.

McFarlane HG, Kusek GK, Yang M, Phoenix JL, Bolivar VJ, Crawley JN. 2008. Autism-like behavioral phenotypes in BTBR T+tf/J mice. Genes, Brain Behav. 7:152–163.

Mike A, Strammer E, Aradi M, Orsi G, Perlaki G, Hajnal A, Sandor J, Banati M, Illes E, Zaitsev A, Herold R, Guttmann CRG, Illes Z. 2013. Disconnection mechanism and regional cortical atrophy contribute to impaired processing of facial expressions and theory of mind in multiple sclerosis: A structural MRI study. PLoS One. 8.

Mori S, Crain BJ, Chacko VP, van Zijl PC. 1999. Three-dimensional tracking of axonal projections in the brain by magnetic resonance imaging. Ann Neurol. 45:265–269.

Mori S, van Zijl PCM. 2002. Fiber tracking: principles and strategies - a technical review. NMR Biomed. 15:468–480.

Mori S, Zhang J. 2006. Principles of Diffusion Tensor Imaging and Its Applications to Basic Neuroscience Research. Neuron. 51:527–539.

Mullen EM. 1995. Mullen Scales of Early Learning, AGS Edition: Manual and Item Administrative Books. Am Guid Serv Inc. 1–92.

Müller RA, Shih P, Keehn B, Deyoe JR, Leyden KM, Shukla DK. 2011. Underconnected, but how? A survey of functional connectivity MRI studies in autism spectrum disorders. Cereb Cortex. 21:2233–2243.

Nordahl CW, Iosif A-M, Young GS, Perry LM, Dougherty R, Lee A, Li D, Buonocore MH, Simon T, Rogers S, Wandell B, Amaral DG. 2015. Sex differences in the corpus callosum in preschool-aged children with autism spectrum disorder. Mol Autism. 6:1–11.

Northam GB, Liegeois F, Tournier JD, Croft LJ, Johns PN, Chong WK, Wyatt JS, Baldeweg T. 2012. Interhemispheric temporal lobe connectivity predicts language impairment in adolescents born preterm. Brain. 135:3781–3798.

Orosco L a., Ross AP, Cates SL, Scott SE, Wu D, Sohn J, Pleasure D, Pleasure SJ, Adamopoulos IE, Zarbalis KS. 2014. Loss of Wdfy3 in mice alters cerebral cortical neurogenesis reflecting aspects of the autism pathology. Nat Commun. 5:4692.

Packer A. 2016. Neocortical neurogenesis and the etiology of autism spectrum disorder. Neurosci Biobehav Rev. 64:185–195.

Pelphrey K a., Morris JP, McCarthy G. 2005. Neural basis of eye gaze processing deficits in autism. Brain. 128:1038–1048.

Pierce K. 2011. Early functional brain development in autism and the promise of sleep fMRI. Brain Res.

Pierce K, Carter C, Weinfeld M, Desmond J, Hazin R, Bjork R, Gallagher N. 2011. Detecting, studying, and treating autism early: The one-year well-baby check-up approach. J Pediatr. 159:458–465.e6.

Pierce K, Marinero S, Hazin R, McKenna B, Barnes CC, Malige A. 2015. Eye-tracking Reveals Abnormal Visual Preference for Geometric Images as an Early Biomarker of an ASD Subtype Associated with Increased Symptom Severity. Biol Psychiatry.

Pinto D, Pagnamenta AT, Klei L, Anney R, Merico D, Regan R, Conroy J, Magalhaes TR, Correia C, Abrahams BS, Almeida J, Bacchelli E, Bader GD, Bailey AJ, Baird G, Battaglia A, Berney T, Bolshakova N, Bölte S, Bolton PF, Bourgeron T, Brennan S, Brian J, Bryson SE, Carson AR, Casallo G, Casey J, Chung BHY, Cochrane L, Corsello C, Crawford EL, Crossett A, Cytrynbaum C, Dawson G, de Jonge M, Delorme R, Drmic I, Duketis E, Duque F, Estes A, Farrar P, Fernandez B a, Folstein SE, Fombonne E, Freitag CM, Gilbert J, Gillberg C, Glessner JT, Goldberg J, Green A, Green J, Guter SJ, Hakonarson H, Heron E a, Hill M, Holt R, Howe JL, Hughes G, Hus V, Igliozzi R, Kim C, Klauck SM, Kolevzon A, Korvatska O, Kustanovich V, Lajonchere CM, Lamb J a, Laskawiec M, Leboyer M, Le Couteur A, Leventhal BL, Lionel AC, Liu X-Q, Lord C, Lotspeich L, Lund SC, Maestrini E, Mahoney W, Mantoulan C, Marshall CR, McConachie H, McDougle CJ, McGrath J, McMahon WM, Merikangas A, Migita O, Minshew NJ, Mirza GK, Munson J, Nelson SF, Noakes C, Noor A, Nygren G, Oliveira G, Papanikolaou K, Parr JR, Parrini B, Paton T, Pickles A, Pilorge M, Piven J, Ponting CP, Posey DJ, Poustka A, Poustka F, Prasad A, Ragoussis J, Renshaw K, Rickaby J, Roberts W, Roeder K, Roge B, Rutter ML, Bierut LJ, Rice JP, Salt J, Sansom K, Sato D, Segurado R, Sequeira AF, Senman L, Shah N, Sheffield VC, Soorya L, Sousa I, Stein O, Sykes N, Stoppioni V, Strawbridge C, Tancredi R, Tansey K, Thiruvahindrapduram B, Thompson AP, Thomson S, Tryfon A, Tsiantis J, Van Engeland H, Vincent JB, Volkmar F, Wallace S, Wang K, Wang Z, Wassink TH, Webber C, Weksberg R, Wing K, Wittemeyer K, Wood S, Wu J, Yaspan BL, Zurawiecki D, Zwaigenbaum L, Buxbaum JD, Cantor RM, Cook EH, Coon H, Cuccaro ML, Devlin B, Ennis S, Gallagher L, Geschwind DH, Gill M, Haines JL, Hallmayer J, Miller J, Monaco AP, Nurnberger JI, Paterson AD, Pericak-Vance M a, Schellenberg GD, Szatmari P, Vicente AM, Vieland VJ, Wijsman EM, Scherer SW, Sutcliffe JS, Betancur C. 2010. Functional impact of global rare copy number variation in autism spectrum disorders. Nature. 466:368–372.

Power JD, Barnes KA, Snyder AZ, Schlaggar BL, Petersen SE. 2012. Spurious but systematic correlations in functional connectivity MRI networks arise from subject motion. Neuroimage. 59:2142–2154.

Pramparo T, Lombardo M V, Campbell K, Barnes CC, Marinero S, Solso S, Young J, Mayo M, Dale A, Ahrens-Barbeau C, Murray SS, Lopez L, Lewis N, Pierce K, Courchesne E. 2015. Cell cycle networks link gene expression dysregulation, mutation, and brain maldevelopment in autistic toddlers. Mol Syst Biol. 11:841.

Redcay E, Courchesne E. 2008. Deviant functional magnetic resonance imaging patterns of brain activity to speech in 2-3-year-old children with autism spectrum disorder. Biol Psychiatry. 64:589–598.

Rohde GK, Barnett a. S, Basser PJ, Marenco S, Pierpaoli C. 2004. Comprehensive Approach for Correction of Motion and Distortion in Diffusion-Weighted MRI. Magn Reson Med. 51:103–114.

Sabers A, Bertelsen FCB, Scheel-Krüger J, Nyengaard JR, Møller A. 2014. Long-term valproic acid exposure increases the number of neocortical neurons in the developing rat brain. A possible new animal model of autism. Neurosci Lett. 580:12–16.

Salvan P, Froudist Walsh S, Allin MPG, Walshe M, Murray RM, Bhattacharyya S, McGuire PK, Williams SCR, Nosarti C. 2014. Road work on memory lane-Functional and structural alterations to the learning and memory circuit in adults born very preterm. Neuroimage. 102:152–161.

Santos M, Uppal N, Butti C, Wicinski B, Schmeidler J, Giannakopoulos P, Heinsen H, Schmitz C, Hof PR. 2011. von Economo neurons in autism: A stereologic study of the frontoinsular cortex in children. Brain Res. 1380:206–217.

Saxe R. 2006. Uniquely human social cognition. Curr Opin Neurobiol. 16:235–239.

Silverman JL, Yang M, Lord C, Crawley JN. 2010. Behavioural phenotyping assays for mouse models of autism. Nat Rev Neurosci. 11:490–502.

Solso S, Xu R, Proudfoot J, Hagler DJ, Campbell K, Venkatraman V, Carter Barnes C, Ahrens-Barbeau C, Pierce K, Dale A, Eyler L, Courchesne E. 2016. DTI provides evidence of possible axonal over-connectivity in frontal lobes in asd toddlers. Biol Psychiatry. 79:676–684.

Sparrow SS. 2011. Vineland adaptive behavior scales. In: Kreutzer JS, DeLuca J, Caplan B, editors. Encyclopedia of Clinical Neuropsychology. ii. ed. New York, NY: Springer New York. p. 2618–2621.

Stoner R, Chow ML, Boyle MP, Sunkin SM, Mouton PR, Roy S, Wynshaw-Boris A, Colamarino SA, Lein ES, Courchesne E. 2014. Patches of disorganization in the neocortex of children with autism. N Engl J Med. 370:1209–1219.

Sugathan A, Biagioli M, Golzio C, Erdin S, Blumenthal I, Manavalan P, Ragavendran A, Brand H, Lucente D, Miles J, Sheridan SD, Stortchevoi A, Kellis M, Haggarty SJ, Katsanis N, Gusella JF, Talkowski ME. 2014. CHD8 regulates neurodevelopmental pathways associated with autism spectrum disorder in neural progenitors. Proc Natl Acad Sci. 111:E4468–E4477.

Symington SH, Paul LK, Symington MF, Ono M, Brown WS. 2010. Social cognition in individuals with agenesis of the corpus callosum. Soc Neurosci. 5:296–308.

Thomas C, Humphreys K, Jung KJ, Minshew N, Behrmann M. 2011. The anatomy of the callosal and visual-association pathways in high-functioning autism: A DTI tractography study. Cortex. 47:863–873.

Travers BG, Adluru N, Ennis C, Tromp DPM, Destiche D, Doran S, Bigler ED, Lange N, Lainhart JE, Alexander AL. 2012. Diffusion Tensor Imaging in Autism Spectrum Disorder: A Review. Autism Res. 5:289–313.

Travers BG, Tromp DPM, Adluru N, Lange N, Destiche D, Ennis C, Nielsen J a, Froehlich AL, Prigge MBD, Fletcher PT, Anderson JS, Zielinski B a, Bigler ED, Lainhart JE, Alexander AL. 2015. Atypical development of white matter microstructure of the corpus callosum in males with autism: a longitudinal investigation. Mol Autism. 6.

Vogan VM, Morgan BR, Leung RC, Anagnostou E, Doyle-Thomas K, Taylor MJ. 2016. Widespread White Matter Differences in Children and Adolescents with Autism Spectrum Disorder. J Autism Dev Disord. 46:1–10.

Wahlsten D, Metten P, Crabbe JC. 2003. Survey of 21 inbred mouse strains in two laboratories reveals that BTBR T/+ tf/tf has severely reduced hippocampal commissure and absent corpus callosum. Brain Res. 971:47–54.

Weinstein M, Ben-Sira L, Levy Y, Zachor D a., Itzhak E Ben, Artzi M, Tarrasch R, Eksteine PM, Hendler T, Bashat D Ben. 2011. Abnormal white matter integrity in young children with autism. Hum Brain Mapp. 32:534–543.

Wolff JJ, Gerig G, Lewis JD, Soda T, Styner M a., Vachet C, Botteron KN, Elison JT, Dager SR, Estes a. M, Hazlett HC, Schultz RT, Zwaigenbaum L, Piven J. 2015. Altered corpus callosum morphology associated with autism over the first 2 years of life. Brain. 1–13.

Wolff JJ, Gu H, Gerig G, Elison JT, Styner M, Gouttard S, Botteron KN, Dager SR, Dawson G, Estes AM, Evans AC, Hazlett HC, Kostopoulos P, McKinstry RC, Paterson SJ, Schultz RT, Zwaigenbaum L, Piven J. 2012. Differences in white matter fiber tract development present from 6 to 24 months in infants with autism. Am J Psychiatry. 169:589–600.

Wolff JJ, Ph D, Gu H, Gerig G, Elison JT, Dawson G, Estes AM, Evans A, Paterson SJ, Schultz RT, Zwaigenbaum L. 2013. NIH Public Access. 169:589–600.

Xiao Z, Qiu T, Ke X, Xiao X, Xiao T, Liang F, Zou B, Huang H, Fang H, Chu K, Zhang J, Liu Y. 2014. Autism spectrum disorder as early neurodevelopmental disorder: evidence from the brain imaging abnormalities in 2-3 years old toddlers. J Autism Dev Disord. 44:1633–1640.

Yeatman JD, Dougherty RF, Myall NJ, Wandell B a., Feldman HM. 2012. Tract Profiles of White Matter Properties: Automating Fiber-Tract Quantification. PLoS One. 7.

Zikopoulos B, Barbas H. 2010. Changes in prefrontal axons may disrupt the network in autism. J Neurosci. 30:14595–14609.

Zilbovicius M, Meresse I, Chabane N, Brunelle F, Samson Y, Boddaert N. 2006. Autism, the superior temporal sulcus and social perception. Trends Neurosci. 29:359–366.

